# A comparison of hard and soft direct methods for DNA extraction from soil

**DOI:** 10.1101/2022.03.07.483395

**Authors:** Patrick Hill, Mathieu F Dextraze, David Kroetsch, Christopher N Boddy

## Abstract

Nucleic acid extraction is the first step in molecular biology studies of soil bacterial communities. The most common used soil DNA extraction method is the direct, hard extraction Mobio method, which uses bead beating to lyse bacteria. In this study we compared the Mobio method with a soft, enzymatic lysis extraction method. Next generation sequencing (Illumina and Pyrosequencing) of amplicons generated from four 16S primer pairs and DNA from 12 soils and 3 composts was used to compare the two extraction methods.

Four bacterial orders, the delta proteobacterial *Desulfuromonadales* and gamma proteobacterial *Pseudomonadales, Enterobacteriales*, and *Alteromonadales* were more common in amplicons from soft extracted DNA, sometimes by two orders of magnitude. These groups can be a significant fraction of the bacterial population. For example the *Pseudomonadales* made up to 16 % and *Enterobacteriales* 10% of amplicons from Soft extracted DNA. The JG30-KF-CM45 order was under extracted by the enzymatic lysis extraction method. Results differed more by primer choice than extraction method and the phylogenetic resolution of differences between extraction methods changed with primer choice.

Given how often Mobio extraction is used, these proteobacterial orders are probably under-represented in the studies of soil bacteria that use nucleic acid methods. Further improvements in soil DNA extraction are needed. Amplicons sequencing studies should use a range of different primers to confirm the phylogenetic resolution of their results.

**Importance:** Several large scale studies of soil bacteria that compare thousands of soil samples across continents have used the Mobio method for DNA extraction. Large scale studies like these are increasing with the recent establishment of the Global Soil Biodiversity Observation Network (Soil BON), which also uses the Mobio method. The results of this work will be used to make policy decisions about how to manage the soil and may be a guide for bioprospectors. As the Mobio method is so widely used, it is important to know its limitations. Studies that use the Mobio method underestimate the fraction of several proteobacterial groups. Most notably the Enterobacteria and Pseudomonas can be under extracted by 10-100 fold. The degree of under extraction varies with different soils.

## Introduction

Most soil bacteria cannot be grown in the laboratory (Torsvik et al., 1990). To study these bacteria without cultivating them, their DNA is extracted from soil and the DNA or PCR products amplified from it are sequenced (Delmont et al., 2011). Bacterial DNA can be extracted from soil indirectly or directly. In indirect extraction, bacteria are taken out of soil before being lysed, in direct extraction bacteria are lysed in place in the soil.

Indirect extraction is used when DNA quality is important. As bacteria are not in the soil when lysed, the extracted DNA is cleaner. Usually “soft” chemical methods such as enzymes and/or detergent are used for lysis so DNA fragments are relatively long (> 50 000 bp). DNA of this length can be useful for several reasons. Longer fragments are more likely to contain the full length functional units, such as natural product biosynthetic gene clusters, which can then be heterologously expressed (Gabor, Alkema, & Janssen, 2004)., As well as this, the longer fragments are, the more easily they can be assembled into genomes. This may be useful as newer long sequencing methods are used in metagenomic sequencing (Haro-Moreno, López-Pérez, & Rodríguez-Valera, 2020)(Goethem et al., 2021) (Waschulin, James, Newsham, Donadio, & Corre, 2022).

Direct extraction is faster and extracts more of the whole bacterial population than indirect extraction (Delmont et al., 2011) and is used in ecological studies that compare the bacterial populations of soils. These extractions typically use “hard” methods that physically lyse bacteria and are faster than soft lysis. The hard lysis shears extracted DNA into short fragments (< 20 000 bp) (Yeates, Gillings, Davison, Altavilla, & Veal, 1998) (Cullen & Hirsch, 1998) (Bremen, Miller, Bryant, Madsen, & Ghiorse, 1999) (Í nceošlu, Hoogwout, Hill, & Van Elsas, 2010). However these studies often use PCR to amplify much shorter lengths (< 1500 bp) of ribosomal genes for sequencing, so DNA length does not affect the results.

Since the early 2000s direct extraction is usually done with kits containing premixed reagents and well tested, standardized methods. Kit extraction usually begins by weighing less than a gram of soil or sediment into a tube with beads. The tube is then shaken for a few minutes so that the beads lyse bacterial membranes and cell walls. An extract containing the DNA of the lysed bacteria is then cleaned and concentrated by precipitating out impurities and binding the DNA to a stationary phase in a column before washing and elution.

The most popular of these kits is the Mobio Power soil kit (modified and renamed the DNeasy PowerSoil Kit). It is used by several large scale projects that compare bacterial populations from thousands of soils across large areas (e.g Australia, (Bissett et al., 2016) the European Union (Orgiazzi, Ballabio, Panagos, Jones, & Fernandez-Ugalde, 2018)) and the Earth Microbiome Project (Thompson et al., 2017). The Mobio Powersoil kit is also the method that will be used in the recently established global Soil Biodiversity Observation Network (Guerra et al., 2021). In this study we refer to this extraction method as the Mobio method.

We compare the Mobio method with a method that is a compromise between direct and indirect methods. It directly extracts DNA from soil using soft lysis (enzymes and SDS). Zhou et al’s (1996) soil DNA extraction method uses enzymes, and detergent to lyse bacteria (Zhou, Bruns, & Tiedje, 1996). When used on clayey bamboo forest soils from the Cauca flood plain near Cali, Colombia, this method yielded thick black liquid that could not be read on spectrophotometers nor amplified with PCR. After several modifications such as washing before extraction and altering extraction and cleaning steps, it yielded long fragments of clean DNA from many soils (Peñ a-Venegas, Cardona, Arguelles, & Arcos, 2007) (Hill et al., 2011) and sediments (Nováková et al., 2005). We call this method the Soft method.

We extracted DNA from soils and compost using both methods, PCR amplified the 16S ribosomal genes using four different primer pairs, and sequenced the amplicons with next generation sequencing. Mobio extracted DNA was more alpha diverse than Soft extracted DNA. However there were often 10-100 × more sequenced amplicons from several gamma proteobacterial orders (*Enterobacterales* and *Pseudomonadales*) in the soft extracted DNA. These bacterial groups may be under-represented in the many studies that use the Mobio method.

## Results and Discussion

This study examines how extraction methods affect our view of soil bacterial populations. Therefore we sampled a broad range of soils only once rather than fewer soils in triplicate as is done to characterize the bacterial community of a particular soil. Twelve contrasting soils and three compost samples were sampled once. DNA was extracted in triplicate from one of these samples (Exp1B) and in duplicate from a second (Exp2A) to make sure that results were reproducible.

Eight soils were from Ottawa, Canada, two were acid forest podzols from the Mattawa plain (Ontario) and Vancouver, Canada and one each was from San Diego, California and North-East Kansas U.S.A. (Table 1). The three compost samples were from the top and bottom of a compost pile receiving garden and household waste (80F and 80F Top) and the top of a compost pile receiving garden waste in a long-term care home (Glebe).

**Table1.**
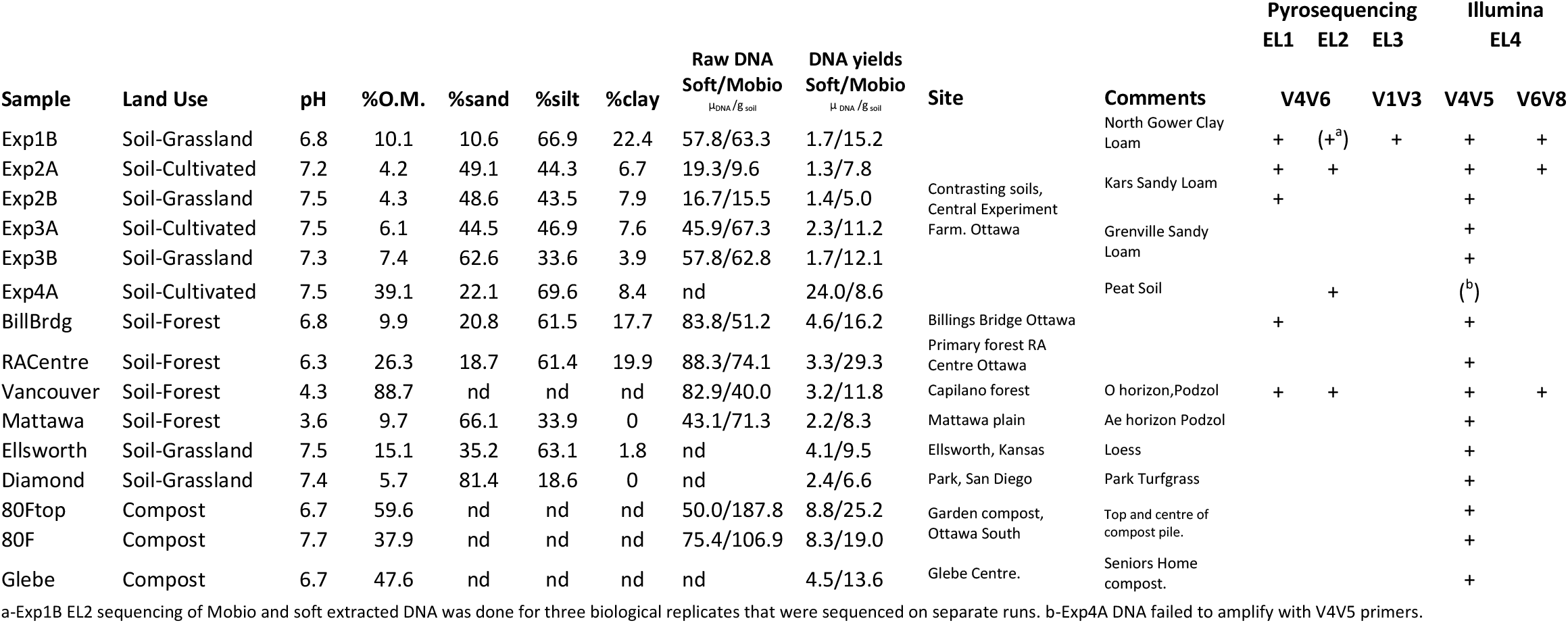
Sample description and amplicon sequencing plan.

DNA was extracted from the samples using the MoBio and Soft DNA extraction methods. The raw uncleaned and cleaned final DNA yields were measured for each extraction method. Four different primers pairs for differing regions of the 16S rRNA gene were used to generate amplicons from the cleaned final DNA samples. Amplicons were sequenced with several methods of next generation sequencing (Table 1).

### Minor effects of extraction method on Alpha and Beta diversity compared to primers and sample

Amplicons from the five soil samples Exp1B, Exp2A, Exp2B, BillingsBridge, and Vancouver were generated with V4V5 and V4V6 primers from DNA extracted using both methods. V4V6 amplicons were sequenced by pyrosequencing and the V4V5 amplicons by Illumina. Amplicons from V4V5 primers were more alpha diverse than those from V4V6 (p<0.002 Chao, p<0.014 Faiths PD Supplemental Table 2). There was no statistically significant effect for extraction method. There was a statistically significant effect of extraction method on alpha diversity on results of Illumina V4V5 amplicon sequences from all 11 soils and 3 composts. Mobio amplicons were more alpha diverse than the Soft extracted amplicons (p<0.013 Chao, Supplemental Table1).

For the five soils with both V4V6 and V4V5 results (Exp1B, Exp2A, Exp2B, BillingsBridge, Vancouver), primers had a statistically significant effect on beta diversity (Figure 1, UniFrac weighted p<0.001, UniFrac Unweighted p<0.001, Bray-Curtis p<0.005 Supplemental Table 3) while extraction method did not. However weighted UniFrac analysis of V4V5 amplicons from seven soils from Ottawa found that extraction affected clustering (p<0.017, Supplemental Table 4).

**Figure 1.**
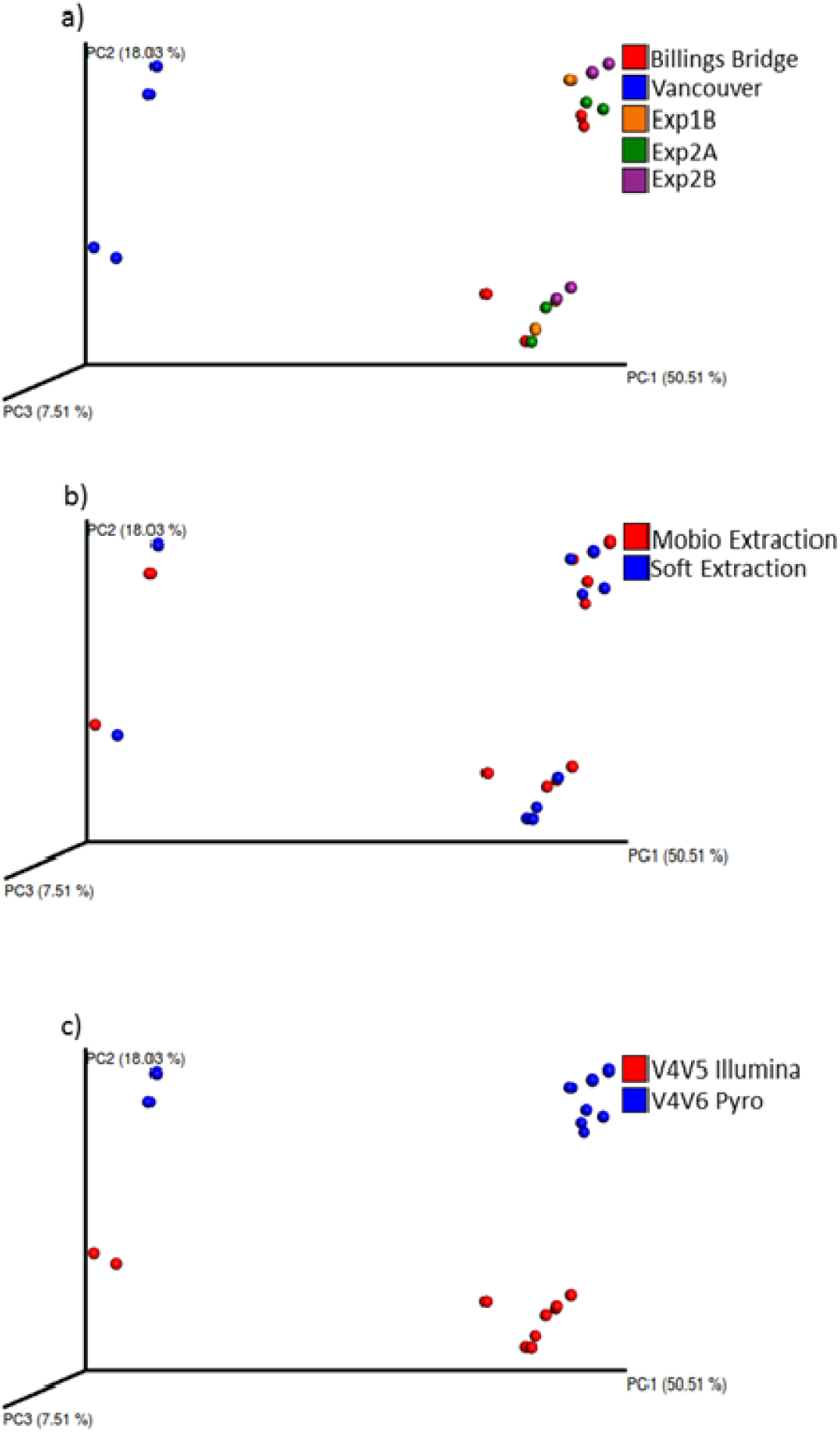
Primers affected the apparent beta diversity of soils more than extraction method. V4V6 Pyrosequencing and V4V5 Illumina amplicon sequencing results for the soils: Vancouver, Billings Bridge, Exp1B, Exp2A, Exp2B. Results are presented through weighted UniFrac Principal co ordinate analysis. Samples are colour coded by a-Soil b-DNA extraction method c-Primer Sequencing method.

There was no evidence that using any combination of primers or extraction method could fundamentally alter understanding of how bacterial communities vary between soils or between soils and other biomes. We compared all of our V4V5 and V4V6 amplicons from soil DNA extracted with both soft and mobio extraction with V4V6 amplicons from non-soil samples: compost, faeces, beach sand, and street dust. UniFrac cluster analysis found that soils clustered together when compared with other environments (Figure2A). There were the same soil outliers using both extraction methods, an urban park soil from San Diego, Diamond and the two acidic podzols (Vancouver, Mattawa). This is not surprising, many previous studies have found that pH strongly affects bacterial populations in soil e.g.(Fierer & Jackson, 2006).

**Figure 2.**
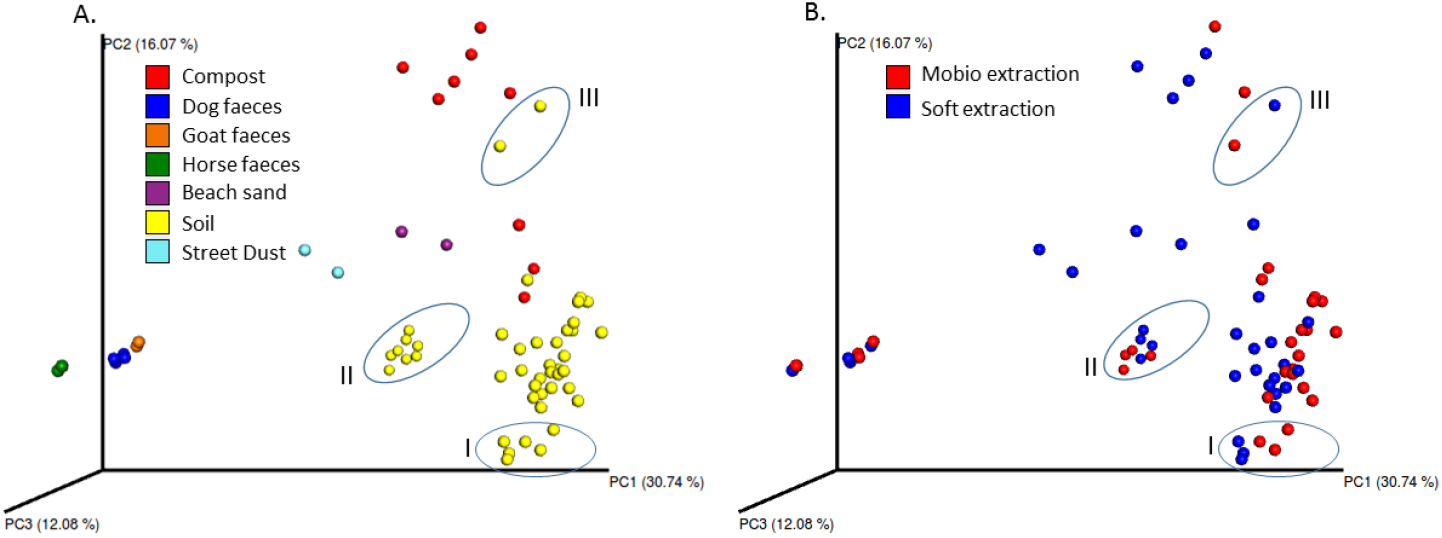
Varying primers and extraction method does not alter overall community differences between soils and other biomes. Pyrosequencing and Illumina sequencing results from this study were combined with pyrosequencing results from other studies from faeces, beach sand and street sediments. Parameters were set to the lowest quality reads in the whole batch (faecal sample amplicons. Minimum length was set at 250 bp and sequences were trimmed with SEED2. Results are presented through weighted UniFrac Principal co ordinate analysis. Samples are colour coded by a-Environment b-DNA extraction method. Three clusters of results are circled. I are the results of V4V6 amplicon pyrosequencing of triplicate samples of DNA from Exp1B extracted with Mobio and soft extraction II are results from two acidic podzols, Mattawa and Vancouver. III are results from Diamond soil from San Diego, USA.

These result match those of earlier comparisons of extraction methods across different biomes and contrasting soils. When several bead beating direct extraction methods were used to compare the bacterial communities of sea water, deep sea sediment, soil, mattress dust, the human mouth, and faeces, samples clustered by environment (Marotz, Amir, Humphrey, Gogul, & Knight, 2017). Similarly, when different direct extraction bead beating methods are used on different soils, samples cluster by soil rather than extraction method (Terrat et al., 2012)(Terrat et al., 2015)(Soliman, Yang, Yamazaki, & Jenke-Kodama, 2017).

However extraction method can affect results if the samples are very similar. For example, when soil samples were taken from the same field at different depths and DNA extracted using several different direct and indirect methods, extraction method strongly affected the results (Delmont et al., 2011).

Similarly 16S DGGE fingerprints from three direct extraction methods that included both soft extraction and Mobio extraction of DNA from three Dutch potato soils showed that the two sandy soils of similar pH clustered by extraction method (Í nceošlu et al., 2010). Lastly, Roldan et al compared the Mobio method to a second direct bead beating method using Illumina 16S amplicon sequencing on three land uses on the same clayey soil and showed that extraction method affected how samples clustered by weighted Unifrac principle coordinate analysis (Roldan, Junca, & Arbeli, 2019).

We found that extraction method was statistically significant on weighted UniFrac clustering of seven Ottawa mineral soils found within five km of each other with a pH range of 6.3-7.5 (Supplemental Table 4). Thus the choice of DNA extraction method may distort results when comparing very similar soils or treatments on the same soil.

### STAMP analysis shows differential extraction of *Enterobacteriales* and *Pseudomonades*

STAMP (Statistical analysis of taxonomic and functional profiles)(Parks, Tyson, Hugenholtz, & Beiko, 2014) group analysis found many statistically significant differences at a range of phylogenetic levels for both extraction method and primers. However while many differences were statistically significant, they were also small and so would not alter the overall view of the bacterial community (e.g. Supplemental Figure S2). When a cutoff for an effect size of at least four-fold change in abundance or greater was used to refine the STAMP output, there were still significant differences for both primers and extraction method.

The largest of these differences was for primers (V4V5 vs V4V6) for the *planctomycete* phylum, which was preferentially amplified by V4V5 primers (Supplemental figure S1). It is uncertain if the V4V6 primers were under amplifying or V4V5 primers over amplifying *planctomycetes*. Mantri et al compared 16S amplicon sequencing and shotgun sequencing of DNA from four forest soils and found that the *planctomycetes* were overrepresented in the amplicons sequences by several orders of magnitude (Mantri et al., 2021), suggesting that some primers can over amplify *planctomycetes*. However this study used V3V4 amplicons and so cannot be directly compared to ours.

Extraction method had smaller effects on the abundance of some phylogenetic groups but this effect was found using several primer pairs and thus is unlikely to be an artifact of amplification. There was no statistically significant difference of extraction method for group level analysis of the five soils with V4V5 and V4V6 amplicon data. There was however for the eleven soils with V4V5 data. Five orders were more common in amplicons from one or the other extraction method (Figure 3). Three gamma proteobacterial orders, the *Pseudomonades, Enterobacteriales, Alteromonadales* and the delta proteobacterial order *Desulfuromonadales* were more abundant in amplicons from Soft extracted DNA. The order JG30-KF-CM45 from the *Thermomicrobia* was more abundant in amplicons from Mobio extracted DNA. (Supplemental tables 2 and 3). The largest effect for differential extraction was observed for the *Pseudomonadales a*nd *Enterobacteriales*. These orders were often a significant fraction of the 16S amplicons. For example the *Pseudomonadales* were 16.5 % of the amplicon from the Soft extraction of Exp2A soil sample and the *Enterobacteriales* were 10.7 % of the amplicon in the Soft extraction of RACentre soil. *Pseudomonadales* and *Enterobacteriales* amplicons were over 100 times more common in Soft extracted than Mobio extracted DNA from Exp1B and Exp2A samples (Supplemental Table 5).

**Figure 3.**
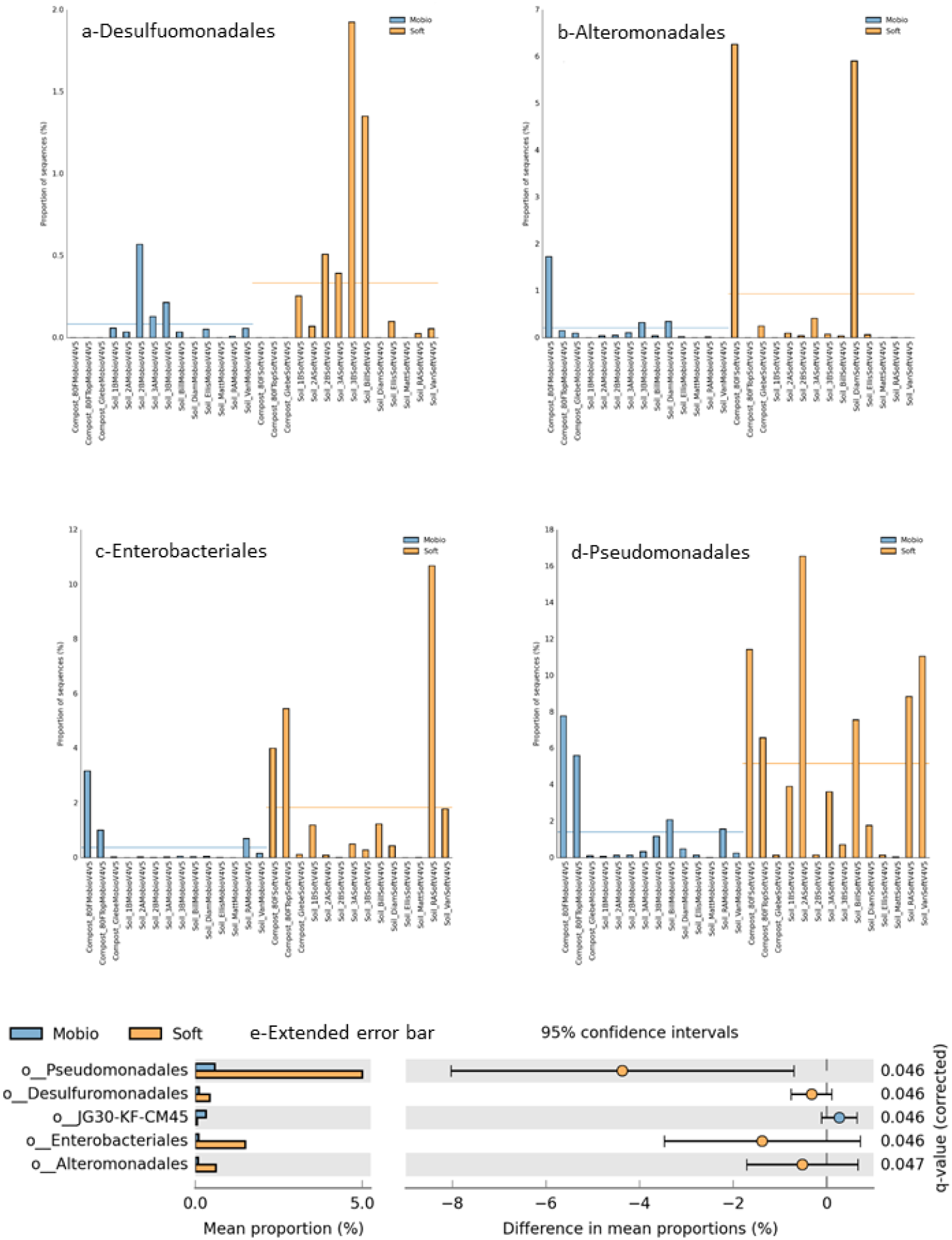
Extraction method affects abundance of different phylogenetic groups from the V4-V5 primer datasets for 11 soils and three compost samples. Bar graphs of the percentages of the orders a*-Pseudomonadales*, b-*Enterobacteriales*, c-*Alternomonadales*, d-*Desulfuromonadales* in soils and compost. e-STAMP Extended error bar plots at the order level for group analysis of soils (Soft versus Mobio extraction). Only results where differences were at least 4 fold and in at least one of the samples the order was more than 1% of sequences are shown.

Differential extraction of *Pseudomonadales* and *Enterobacteriales* was confirmed with a range of amplicon sequencing methods. All of the primer pairs used found preferential extraction of the *Enterobacteriales* with the Soft extraction method, but the genera amplified changed with primer and PCR conditions. V4V5 and V6V8 Enterobacterial amplicons were overwhelmingly unclassified *Enterobacteriaceae*. V4V6 Enterobacterial amplicons sequenced by EL1 were often *Pantoea* while the V4V6 amplicons sequenced by EL2 were often *Yersinia*. This suggests that Soft extraction preferentially extracts a broad range of *Enterobacteriales*. Similar Pseudomonal genera were also sequenced from amplicons generated from all primers except the V4V6 primers (Supplemental Tables 5). An example of the differences in results with different primers is shown in figure 4 for Mobio and Soft extracted DNA for sample Exp1B.

**Figure 4.**
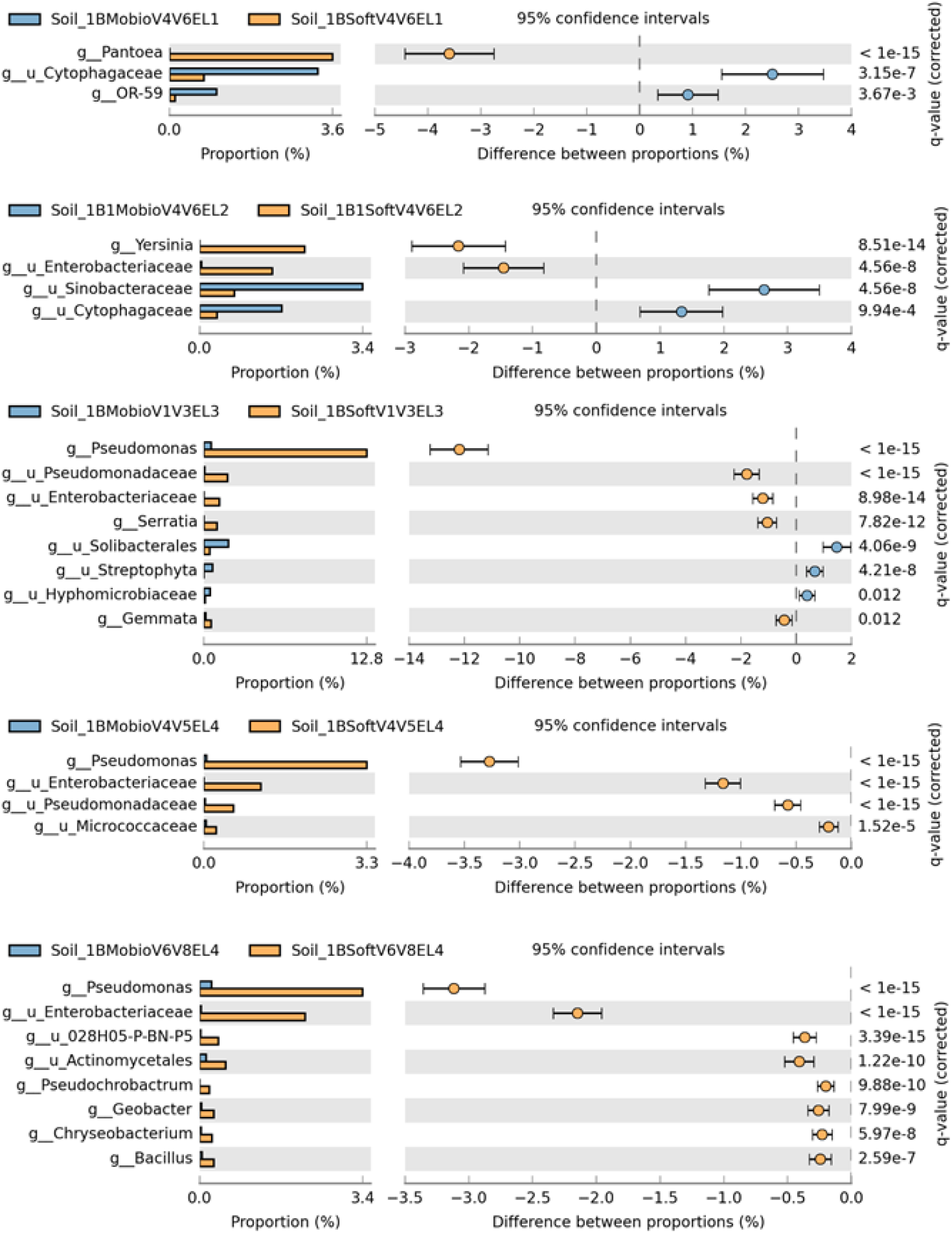
Comparisons of extraction method give different results with different primers and PCR conditions. Examples of STAMP Extended error bar plots at the genus level of soil Exp1B comparing extraction methods from four external sequencing laboratories and primer pairs. Results are from a-EL1/V4V6 primers, b-EL2/V4V6 primers, c-EL3/V1V3 primers, d-EL4/V4V5 primers, e EL4/V6V8 primers. Only results that are four-fold or more different between extraction method are shown. Multiple test correction with Storey FDR.

Other bacterial groups were under-extracted by the Mobio method although results were only found for one or two samples with a single set of primers. V4V6 amplicon sequencing found that amplicons from the *Bacillales* order and the genus *Stenotrophomonas* (a gamma proteobacterial group in the *Xanthomonadales* order), were overwhelmingly more common in Soft extracted DNA from two soils (Figure 5, Supplemental Table 6,7) although the *Bacillales* were overall more common in Mobio extracted DNA (Supplemental Figure 2). The largest fraction of the bacterial community preferentially extracted by either method in any one sample was the *Streptophytes* (from Chloroplasts) by Soft extraction in 80F top compost sample. These made up 35% of V4V5 amplicons from Soft extracted DNA, 261 fold more than Mobio DNA. (Figure 6, Supplemental Table 7).

**Figure 5.**
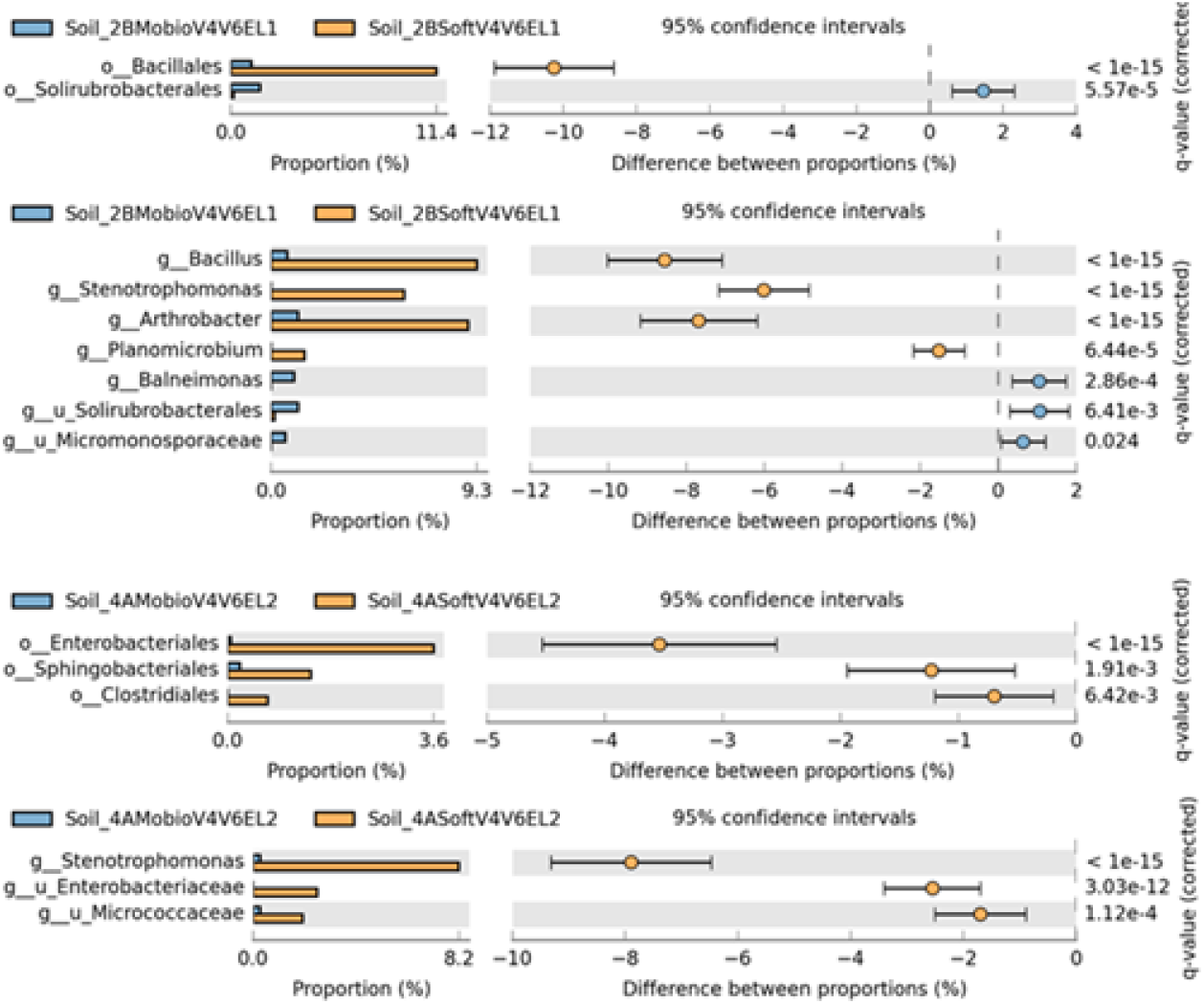
The *Bacillales* order and *Stenotrophomonas* genus were preferentially extracted by Soft extraction in two soils. STAMP Extended error bar plots at the genus and order levels of soils Exp2B and Exp4A. Results are from two external sequencing laboratories, using the V4-V6 primers (EL1, EL2), EL1 for Ex2B, EL2 for Exp4A. Sequences from EL2 and EL3 were trimmed to 450bp. Only results that are four-fold or more shown. Multiple test correction with Storey FDR.

**Figure 6.**
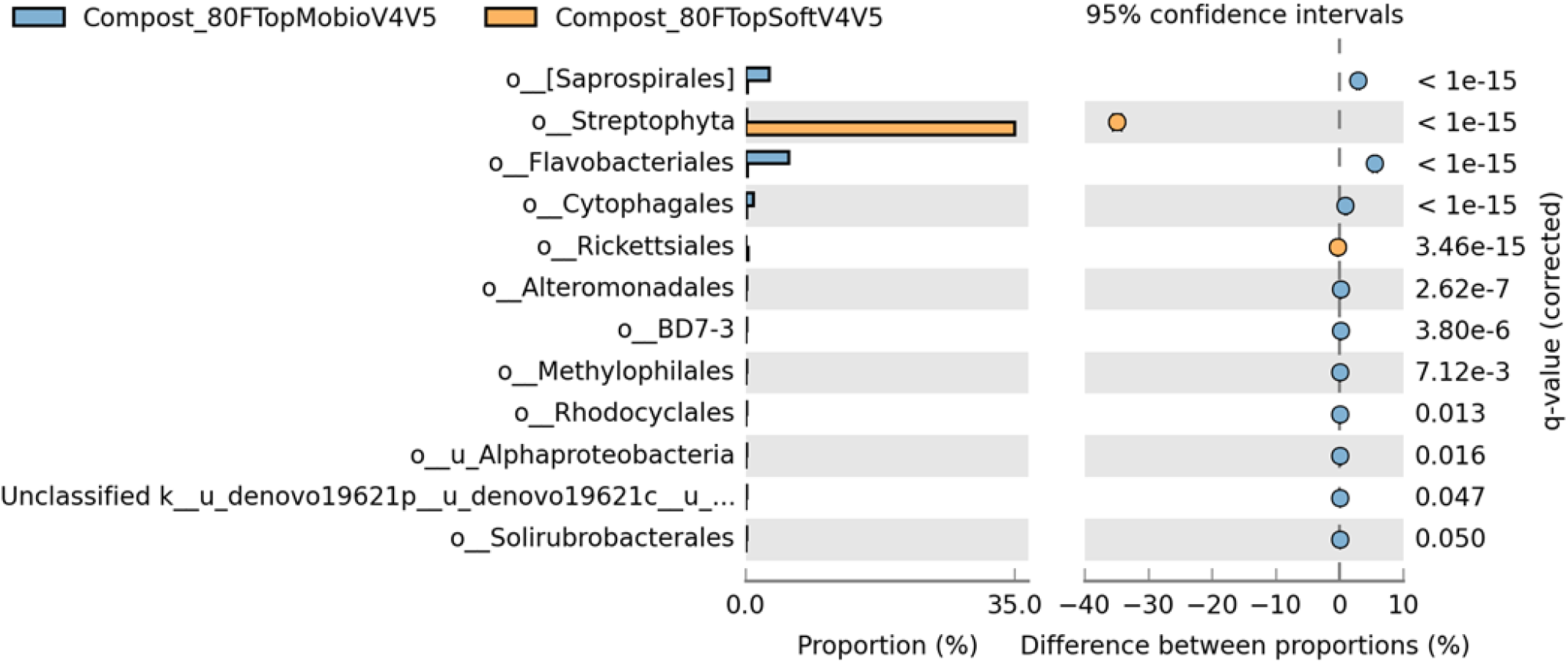
Preferential extraction of the *Streptophyta* order in compost. STAMP Extended error bar plots at the order level from the top layer of garden compost 80F. Results are from EL4, using the V4-V5 primers. Only results that are ten-fold or more shown. Multiple test correction with Storey FDR.

### Extraction efficiency

We found that the amplicons from several groups were much more abundant in PCR products from soft extracted DNA than Mobio extracted DNA. A possible reason for this is that the Soft extraction method did not extract more of these groups but rather a lot less of everything else. For this to be true, the Soft extraction would have to extract tens to hundreds times less DNA than the Mobio method from the whole bacterial population in some soils and similar amounts in others. We tested this by measuring how much DNA was extracted by each method in eleven of our samples. Most of the DNA extracted from soil is bacterial (Fierer et al., 2012) (Frisli, Haverkamp, Jakobsen, Stenseth, & Rudi, 2013) (Kerfahi et al., 2019) so the DNA yield before any is lost during cleaning (Raw yield) is a reasonable estimate of how well the bacterial community is sampled.

Raw DNA yields were similar for both extraction methods (Table 1) and comparable to raw DNA yield from soils in other studies (Zhou et al., 1996)(Bürgmann, Pesaro, Widmer, & Zeyer, 2001) (Carrigg, Rice, Kavanagh, Collins, & O’Flaherty, 2007) (Marstorp & Witter, 1999) (Taylor, Wilson, Mills, & Burns, 2002)(Yokoyama et al., 2017). Thus our STAMP results were not caused by the Soft method under extracting DNA.

DNA extraction efficiency is often measured by apparent alpha diversity of the sample, most simply measured as species richness (how many species are extracted) (Í nceošlu et al., 2010) (Terrat et al., 2012) (Hermans, Buckley, & Lear, 2018) (Zielińska et al., 2017). Efficiency can also be measured by how well the method extracts from “mock communities” of known mixtures of different bacteria (Hallmaier-Wacker, Lueert, Roos, & Knauf, 2018) (Hermans et al., 2018) (Morgan, Darling, & Eisen, 2010).

Both of these strategies are a good way of comparing extraction methods if what limits extraction is the diversity of cell walls and membranes of bacteria. Some of our results suggest that this is so. Bead beating is better than enzymatic extraction in lysing bacterial spores and we found that the Mobio kit extracted more of the *Actinomycetales* and *Bacillales* orders than Soft extraction. This was also the case for the JG30-KF-CM45 order and Alpha proteobacteria (Supplemental Table 2).

Others results from our study do not match this assumption. It is unlikely that Soft extraction worked better than Mobio extraction on the cell walls and membranes of such different groups as the gammaproteobacterial Pseudomonadales, Enterobacteriales, and Alteromonadales, and the deltaproteobacterial Desulfuromonadales. As well, if the differences in the ability of the Mobio and soft extraction methods to degrade cell walls and membranes were responsible for these differences, the ratio for each sample should be similar. Instead they varied by orders of magnitude. For example with the Ilumina V4V5 Pseudomonadales amplicons the Soft/Mobio ratio ranges from 118.3 in Exp2A to 0.6 in Exp3B (Supplemental Tables 5, 6, 7.).

DNA extraction from soil may be limited by the soil itself rather than the membranes and cell walls of soil bacteria. Our Soft lysis method is unusual in that it includes a washing step with Tris and EDTA. Washing soil before cell lysis has several effects. It reduces extraction of humic materials, adjusts pH, and removes contaminants all of which help lytic enzymes work. (LaMontagne, Michel, Holden, & Reddy, 2002) (Fortin, Beaumier, Lee, & Greer, 2004) (Stoeva et al., 2014). Washing may also remove relic DNA, the DNA remaining in soil after cell death. Removing relic DNA can reduce DNA yield and apparent alpha bacterial diversity (Carini et al., 2016)(Carini et al., 2020). This may partly explain the lower apparent alpha diversity and some lower DNA yields for soft extracted DNA.

In contrast, He *et al* found that soil washing increased DNA yield (He, Xu, & Hughes, 2005). He *et al* ascribe this to washing reducing DNA adsorbtion to clay and increasing soil dispersion. While DNA adsorbtion is unlikely to cause preferential extraction of bacterial orders, soil dispersion might.

Soil dispersion is the breaking up soil aggregates into their individual sand, silt and clay particles. Large aggregates (> 200 μm), are easily disrupted. Smaller aggregates are more resistant to physical disruption. Aggregates between 2 and 20 μm can be very resistant to physical disruption and are often made of bacteria surrounded by polysaccharides and clay particles (Tisdall & Oades, 1982). These bacterial microaggregates are bound together by surface charges and cation bridging between organic matter and clay particles and can be very stable, lasting even after bacteria die (Totsche et al., 2018). As they are resistant to mechanical degradation, the microaggregates surrounding bacteria may protect them from Mobio bead beating. Washing with EDTA could remove the stabilizing cations, releasing bacteria from the microaggregates. This could explain the preferential extraction of *Pseudomonadales* and *Enterobacteriale*s by soft extraction.

As seen in our results, different soils would release different numbers of bacteria. Not all soils have as many or as stable aggregates. Extremely sandy soils (Arenosols) are structureless and aggregation is depends on clay content and mineralogy (Totsche et al., 2018).

As also seen in our results, the released bacteria would also be from different groups than those of the bulk soil. The bacterial populations in small aggregates differ from the community in the whole soil (Ranjard & Poly, 2000) (Sessitsch, Weilharter, Gerzabek, Kirchmann, & Kandeler, 2001)(Mummey, Holben, Six, & Stahl, 2006). Three recent studies have used bead beating DNA extraction and next generation sequencing of 16S amplicons to characterize the bacterial communities of soil aggregates to phylogenetic levels at or below the order level (Bach, Williams, Hargreaves, Yang, & Hofmockel, 2018)(Fox et al., 2018)(Biesgen, Frindte, Maarastawi, & Knief, 2020). All found differences in statistically significant differences in bacterial populations of microaggregates compared to other aggregate sizes and the bulk soil.

## Conclusion

This study compared two very different direct soil DNA extraction methods. The Mobio method is fast, convenient, and scalable and like most methods used in Microbial Ecology, uses bead beating to lyse bacteria. The Soft method is much slower and less convenient as it is uses enzymes and detergents for lysis. Results from both methods gave the same broad view of how soil bacterial populations differ from other environments and each other. We show that the soil extraction methods can under extract certain groups of bacteria and thus misrepresent how similar soils differ from each other but do not distort the overall picture of soil bacterial diversity.

We found that that primers and PCR conditions can affected 16S amplicon sequencing results (Figure4 Supplemental Tables 5,6,7), as have many other earlier studies (Wu et al., 2010) (Ahn, Kim, Song, & Weon, 2012) (Rintala et al., 2017) (Hallmaier-Wacker et al., 2018)(Sze & Schloss, 2019) (Lerma et al., 2020). We also observed that results for different primers all confirmed that the soft method preferentially extracted Enterobacteria, even though they amplified different Enterobacterial genera. Based on this, we suggest that important results be confirmed with a range of PCR primers.

Extraction method biases may be less than primer/PCR biases but may be more important. Many different primers and PCR conditions are used in amplicon sequencing studies, which limit the potential for systemic bias. In contract only a few extraction methods are used. Almost all of these method use bead beating methods similar to the Mobio method to extract DNA. Any biases common to bead beating extraction may cause systematic biases across the scientific literature. Furthermore, as direct sequencing of soil DNA is used, primer biases will become less important while extraction biases will remain.

We show that the Mobio method under-extracts several bacterial orders, particularly the *Pseudomonadales* and *Enterobacteriale*s, sometimes by orders of magnitude as compared to the Soft extraction method. As well as this, the degree of under-extraction varies from soil to soil. This suggests that these groups may be systematically under-represented in the literature. They also may be thought to be common or rare in soils where they are easier or more difficult to extract. When different studies or results from large scale studies are compared this could give a false picture of their distribution.

Under-extraction of the *Pseudomonadales* may mean that some of the secondary metabolites in soil are missed by metagenomics studies. Pseudomonads are one of most common bacterial producers of antibiotics (Bérdy, 2005) and secondary metabolite production by *Pseudomonadales* is important in soil suppressivity (Haas & Défago, 2005), the ability of the soil microbial community to prevent plant diseases.

## Methods

### Sample sites

Twelve soils were sampled. Seven were from Ottawa, Canada, two were acid forest podzols from the Mattawa plain (Ontario) and Vancouver, Canada and one each was from San Diego, California and North-East Kansas U.S.A. (Table 1). The three compost samples were also sampled. They were from the top and bottom of a compost pile receiving garden and household waste (80F and 80F Top) and the top of a compost pile receiving garden waste in an old peoples home (Glebe).

### Sample collection and storage

Soil samples were taken by driving a metal core 5 cm into the surface of the soil five to seven times in a meter square area. For forest soils organic litter was cleared to get to the surface of the mineral layer of soil, for the Mattawa soil the mineral soil was dug away until the Ae horizon could be seen. Compost samples were taken from smaller areas on the scale of centimetres. Samples were mixed by hand in a plastic bag and a two to three gram sub-sample was taken. Samples for the two extraction methods (Mobio/Soft extraction) were taken from this sub-sample. Samples were frozen at −20 °C > 12 hours before extraction.

### DNA Extraction

DNA was extracted using the Mobio and soft extraction method once for 10 of the soils and all three compost samples. For two samples DNA was extracted either twice (Exp2A) and three times (Exp1B) using both methods.

### Mobio DNA extraction

This method is similar to the Earth Microbiomes version of the Mobio extraction method (as the manufacturer’s instructions but with an initial 10 min incubation at 65 °C). We used bead beating tubes rather than 96 well plates that the Earth Microbiome does (http://www.earthmicrobiome.org/emp-standard-protocols/dna-extraction-protocol/).

0.25 g of sample was weighed into plastic tubes with glass beads from the Mobio Powersoil kit, the tubes were incubated in a water bath for 10 min at 65 °C and then shaken with the Mobio adaptor fitted onto a Scientific Industries Vortex-Genie 2 benchtop vortexer. All steps after this were as in the manual provided with the Mobio Powersoil kit.

As the enzymatic extraction method extracted 10-100 times more gamma proteobacterial DNA than the Mobio method from several soils, bead beating was intensified for Ottawa sample Exp1B and Exp2A. The bead beating time was doubled from 10 to 20 min (1BExp20mins, 2AExp20mins). Instead of using a benchtop vortex a more powerful Retsch GmbH - Mixer Mill MM 301 grinder was used for three 30 s periods at maximum speed and the tube was cooled on ice water between bead beating (1BExp30sec) DNA.

### Soft DNA extraction

This method uses enzymatic and SDS lysis. As several previous descriptions of this method were incomplete, it is described in full detail in the supplementary material (Supplemental Data1-Methods).

1.0-2.0 g of sample is weighed into a 50 mL Falcon tube. 45 mL of washing buffer (50 mM EDTA/50 mM Tris/HCl pH 8.3) is added and tubes are then centrifuged at 2643 × g (3500 rpm on a Sorvall Legend RT+ centrifuge) at 4 °C for 60 min. The supernatant is poured off, before storing at −20 °C for > 12 hours.

To begin extraction 2.5 mL of enzymatic extraction buffer (500 mM NaCl, 50 mM Tris, 50 mM EDTA, pH 8.0) is added to pellets for digestion with 12.5 mg of lysozyme. Tubes are incubated for 1 h. 140 μL of 20% SDS, is then added and mixed before adding 1 mg of proteinase K for a further hour of digestion.

Five millilitres of SDS extraction buffer (500 mM NaCl, 300 mM succinic acid, 10 mM ETDA, pH 5.7) is then added, followed by 700 μL of 20% SDS before a 45-min incubation at 65 °C. This buffer is added to lower the pH and raise the salinity, reducing the humic material that goes into solution at 65 °C.

Samples are centrifuged at 2643 × g for an hour at 4 °C. 8 mL of the supernatant is transferred to a 15 mL falcon tube. 1 mL of NaCl (5M) is added to each tube and mixed before 1 mL of 10% cetyl trimethyl ammonium bromide (CTAB) is added and incubated for 30 min at 65 °C. The tubes are then cooled to 4 °C on ice and 8 mL of chloroform is added. The tubes are gently mixed and centrifuged at 4000 × g in a centrifuge at 4 °C. The supernatant is removed. A half volume of 0.5 by weight PEG 6000 in water is added to the supernatant, mixed and left overnight at 4 °C in a 15 mL falcon tube to precipitate the DNA.

Samples are transferred to a 50 mL falcon tube, centrifuged at 4500 × g at 4 °C for 3 hours and the supernatant discarded. The pellets are washed with cold 70% ethanol before being dissolved in 400 µL of TE and transferred to a 1.5 mL Eppendorf tube. The samples are then cleaned with an equal volume of phenol/chloroform/isoamylalcohol (25:24:1) followed by one volume of chloroform/isoamylalcohol (24:1) at 4 °C.

5 µL of CaCl_2_ (0.57 M) is then added for each 100 µL of the supernatant left over after previous steps and the solution is mixed and incubated at 65 °C for 30 min. The tubes are then centrifuged at 14 000 ×g in a bench top mini centrifuge for 10 min.

Finally the supernatant from the CaCl_2_ cleaning is transferred to a new Eppendorf tube and DNA precipitated with 1/10 volume of sodium acetate (3 M, pH 5.7) and 1 volume of isopropanol at room temperature for 10 min. The tubes are then centrifuged at 14,000 × g with a bench top mini centrifuge and the supernatant removed. Pellets are then washed with cold 70% ethanol and dissolved in 50 µL of TE.

### Measuring DNA concentration

Final DNA concentrations were measured using the PicoGreen double stranded DNA dye and the Tecan infinite F200 fluorescence microplate reader with Fluorescence Top Reading and excitation 485 nm/ emission 535 nm. To measure DNA concentration, 1-2 µL of the final DNA sample was brought to 100 µL with TE and mixed with 100 µL of 1:200 dilution of PicoGreen dye in TE on a 96 well plate and read on the plate reader. Standards between 25-1000 ng/mL of λ DNA were used to generate a standard curve for quantification.

For all but four samples (Glebe compost and the soils Exp4A, Ellsworth, Diamond) the DNA concentration was also measured immediately after the final lysis step (Mobio bead beating or the Soft hot SDS pH 5.7 step). To prevent interference, serial dilutions, which gave the equivalent of 0.02-0.10 µL of lysate for were used. Standards were between 0.025-25 ng/mL.

### Sequencing overview

Samples were sent to four different external laboratories (EL) for pyrosequencing or Illumina sequencing. Pyrosequencing was done at EL1 and EL2 for the V4/V6 regions and EL3 for the V1/V3 regions. lllumina sequencing for the V4/V5 and V6/V9 regions was done at EL4. A summary of sequencing is shown in table 1.

Pyrosequencing results (V4/V6 EL1) from Soft extracted DNA from beach sand, faeces and street dust samples are also included in Figure 2. These samples were included to show the relative effect of the two extraction methods and primers on Principal Coordinate Analysis compared to the differences between completely different environments.

### Pyrosequencing

Three versions of pyrosequencing were carried out, two with primers for the V4 to V6 variable regions of the 16S ribosomal subunit, a third with primers for the V1 to V3 variable region.

#### V4-V6 pyrosequencing (EL1 and EL2)

25-50 ng of each sample DNA was used as a template for three independent PCR reactions in 25 µL PCR reaction. The 16S Eubacterial primers 530F (GTG CCA GCM GCN GCG G) and 1100R (GGG TTN CGN TCG TTG) were at 0.5 µM concentrations. The reaction cocktail also included 10x New England Buffer, 0.16 mM Bovine Serine Albumen (New England), 0.5 mM d NTP and 1.25 U of New England Taq.

Cycling steps were 95 °C for 2 min followed by 25 cycles of 95 °C for 1 min; 60 °C for 45 s and 68 °C for 45 s and a final elongation step at 68° C for 5 min.

PCR products were either then sent to an external laboratory for bar coded PCR (EL2) or amplified in house (EL1).

If amplified in house, three independent reactions were again run for each PCR. PCR template was 1 µL of the first PCR, with reaction conditions as before but without Bovine Serine Albumen and for only ten cycles. Bar coded 530-F primers were used in the place of 530-F. PCR product concentration for each sample was measured with Pico Green and the samples combined in equal amounts for a single batch of pyrosequencing. These samples were sent for sequencing on an eighth of a pyrosequencing PCR plate to External Laboratory 1 (EL1).

If sent to external laboratory 2 (EL2) for bar coded PCR, cycling conditions were 94 °C for 3 min followed by 28 cycles of 94 °C for 30 s, 53 °C for 40 s and 72 °C for 1 min and a final elongation step of 72 °C for 5 min. The HotStarTaq Plus Master Mix Kit (Qiagen, Valencia, CA) was used in PCR reactions. PCR products from different studies were combined at this laboratory. Samples sent to EL2, were processed in three different runs, each including an Exp1B sample.

#### V1-V3 pyrosequencing (EL3)

Mobio and Soft extracted DNA from 1BExp was sent to a third sequencing laboratory (EL3) for pyrosequencing the V1 to V3 region, with the 27-F (CAC ATG TGA CGA GTT TGA TCM TGG CTCA G) and 518-R primers (CAC ATG TGA CGA GTT TGA TCM TGG CTC AG).

### IIlumina Sequencing

Samples were sent to fourth external sequencing laboratory (EL4) for Illumina sequencing. There a single PCR amplification was carried out with either the 515-F (GTG YCA GCM GCC GCG GTA A) and 962-R (CCG YCA ATT YMT TTR AGT TT) amplicons (V4-V5 sequencing) or the B969-F (ACG CGH NRA ACC TTA CC) and BA1406-R (ACG GGC RGT GWG TRCA A) amplicons (V6-V8 sequencing) using labelled primers for Mi Seq paired end sequencing.

### Sequence Processing

All sequences were edited for quality and length using the Initial processing step of the Ribosomal Database Program (Cole et al., 2014). Pyrosequencing Q values were set at 25 for soil results from EL1 and EL3 or 20 for all other results. Illumina sequencing Q values were set at 27.

Sequences from EL1 included the full length between the 530-F and 1100-R primers and so both primers were used for processing. Sequences from EL2 and EL3 were shorter, so only the forward primer was used and minimum and maximum lengths were set at 450 and 600 bp. These sequences were then trimmed to the minimum 450 bp length using the SEED2 program (Větrovský, Baldrian, & Morais, 2018). When sequences from EL1 and EL2 were analysed together, all sequences were trimmed to 450 bp.

After this initial processing and trimming, all sequences were aligned with the PyNAST program using the default reference alignment in Qiime1. PyNAST rejects sequences that do not align with the reference alignment. These sequences were usually from non-ribosomal genes (Caporaso et al., 2010). The output of PyNAST was then aligned using the Ribosomal Database Program to check that all sequences aligned to the correct region of the 16S subunit. Sequences were then chimera cleaned with UCHIME in the Ribosomal Database (Edgar, Haas, Clemente, Quince, & Knight, 2011).

Further processing was done with Qiime1. Operational taxonomic units were picked de novo with the pick_otus.py (Edgar, 2010), their representative sequences with pick_rep_set.py and their taxonomy assigned with assign_taxonomy.py (DeSantis et al., 2006). All three of these scripts were run with default values. For the next step align_seqs.py, alignment was with MUSCLE (Edgar, 2004). After filtering (filter_alignment.py) sequences were treed with fasttree (Price, Dehal, & Arkin, 2010) and an OTU table made (make_otu_table.py).

### Alpha and Beta diversity analysis with Qiime

Rarefaction curves were made with the alpha_rarefaction.py script. Alpha rarefaction was compared with using the multiple_rarefactions.py, alpha_diversity.py, collate_alpha.py and compare_alphadiverisity.py scripts. Beta diversity was measured using the beta_diversity_through_plots.py script to run unweighted and weighted UniFrac principal coordinate analysis (Lozupone & Knight, 2005). EMPeror and jackknifed UPGMA clustering was used to present the results of this work (Vázquez-Baeza, Pirrung, Gonzalez, & Knight, 2013). To test if the differences that were seen on the PCoA plots and UPGMA trees was statistically significant, Anosim analysis was run using the script compare_categories.py.

### 16S library comparison with Library compare/STAMP

The 16S sequences that were extracted from using each method for each sample were compared pairwise with the Library Compare function of the Ribosomal Database (Cole et al., 2014). Library Compare compares two bacterial communities across the phylum to genus levels of phylogeny however the significance of each difference is not corrected for multiple tests (if enough phylogenetic levels are tested some will randomly be significant) or sample effect (differences may be statistically significant but small).

STAMP (Statistical analysis of taxonomic and functional profiles) is a program that only compares bacterial populations at a single phylogenetic level at a time, but it corrects results for multiple tests and sample effect (Parks et al., 2014). STAMP was used to confirm and present the results of Library Compare.

To make STAMP files the Qiime1 representative sequence file was assigned taxonomy with uclust. The assigned taxonomy file was then reformatted with a python script. This script fills in missing column entries where there is no taxonomic information (SupplementalDataMethods-2). This reformatting is necessary for further processing of the data. The modified taxonomy file and a list of operation taxonomic units (from the pick_otu.py script) was used to make a biom file (make_otu_table.py). The biom file was then converted to json format and used in STAMP to create a STAMP (.spf) file. The species and OTU columns were removed in the LibreOffice Calc spreadsheet.

This spf file was then checked with the check hierarchy script available at http://kiwi.cs.dal.ca/Software/STAMP, and corrected in LibreOffice Calc before being uploaded into STAMP with a mapping file.

Sequence results from each laboratory were compared using two group analysis (Mobio vs Soft extraction) and laboratory results from each sample using different extraction method were compared using two sample comparison. Comparisons were carried out with the recommended settings, for two group analysis Welch’s t-test/Welch’s inverted/Storey’s FDR, for two sample analysis Fisher’s exact test/DP: Newcombe-Wilson/Storey’s FDR. Filtres were set with a p value of 0.05 and to identify groups where differences between extraction methods were not only statistically significant but large. For two group analysis this was with an effect size for rations of 4 (differences less than 4 × would not be reported) for two sample analysis the effect size for ratios set at 10. A maximum value screen of 240 was set for Illumina data as they had many more sequences (at least one samples of the must have at least 240 sequences from the group).

### Accession Numbers

Raw sequence data for this paper can be found in the NCBI SRA database as BioProject PRJNA482847.

## Acknowledgements

The broad sampling that this study used means that we owe thanks to many who helped us. The beach sample from Cornwallis Island was collected by Debbie Iqaluk and Alexandre Poulain; the sampling was made possible thanks to an NSERC discovery grant awarded to Alexandre Poulain, a Research license from the Nunavut Research Institute (license 02-109-11R), the logistical support of PCSP and the consent of the Resolute Bay Hunter’s and Trapper’s Association. Dr. Rashid Nazir provided the Faisalabad Clock tower street dust sample, Panos Argyropoulos the Vancouver forest soil sample, Keavin M. Stanford-Finnerty the beach sample from Prince Edward Island. An employee of the Glebe Centre showed us a compost heap that was in use in a long-term care facility. Douglas Millar set up collaboration with Kara-Lee Golota, allowing us to take dung samples at her farm the Gathering Place Sanctuary. Matt Dextraze was supported by an Ontario Graduate Scholarship. Texture analysis of samples was done with the help of Jean Bjornson of the Geography Dept of the University of Ottawa. Fred Meyer provided the 80F and 80F Top compost samples, was interested in the work, but did not live to see the results.

## Contributions

D.K. sampled Ottawa soils and advised on sampling strategy, P.H. performed the laboratory work, data analysis and wrote the first version of the manuscript, M.F.D. wrote and ran the script that allows Qiime1 taxonomic data to be used in STAMP, C.N.B. contributed to experimental design, critically reviewed the manuscript and sampled the Mattawa, Ellsworth and Diamond samples.

The manuscript was reviewed and approved by all authors.

